# cpSRP43 is both highly flexible and stable: Structural insights using a combined experimental and computational approach

**DOI:** 10.1101/2022.01.11.475959

**Authors:** Mitchell Benton, Mercede Furr, Vivek Govind Kumar, Feng Gao, Colin David Heyes, Thallapuranam Krishnaswamy Suresh Kumar, Mahmoud Moradi

## Abstract

The novel multidomain protein, cpSRP43, is a unique subunit of the post-translational chloroplast signal recognition particle (cpSRP) targeting pathway in higher plants. The cpSRP pathway is responsible for targeting and insertion of light-harvesting chlorophyll a/b binding proteins (LHCPs) to the thylakoid membrane. Nuclear-encoded LHCPs are synthesized in the cytoplasm then imported into the chloroplast. Upon emergence into the stroma, LHCPs form a soluble transit complex with the cpSRP heterodimer, which is composed of cpSRP43 and cpSRP54, a 54 kDa subunit homologous to the universally conserved GTPase in cytosolic SRP pathways. cpSRP43 is irreplaceable as a chaperone to LHCPs in their translocation to the thylakoid membrane and remarkable in its ability to dissolve aggregates of LHCPs without the need for external energy input. In previous studies, cpSRP43 has demonstrated significant flexibility and interdomain dynamics. However, the high flexibility and structural dynamics of cpSRP43 is yet unexplained by current crystal structures of cpSRP43. This is due, in part, to the fact that free full length cpSRP43 is so flexible that it is unable to crystalize. In this study, we explore the structural stability of cpSRP43 under different conditions using various biophysical techniques and find that this protein is concurrently highly stable and flexible. This conclusion is interesting considering that stable proteins typically possess a non-dynamic structure. Molecular dynamics (MD) simulations which correlated with data from biophysical experimentation were used to explain the basis of the extraordinary stability of cpSRP43. This combination of biophysical data and microsecond-level MD simulations allows us to obtain a detailed perspective of the conformational landscape of these proteins.

## Introduction

The signal recognition particle (SRP), SRP receptor and translocase are components of a conserved and ubiquitous protein targeting system (Gutensohn et al. 2006). The SRP pathway, mediated by the ribosome, co-translationally targets proteins to the endoplasmic reticulum (ER) of eukaryotes and the inner membrane of prokaryotes (Li et al. 1995). The chloroplast signal recognition particle (cpSRP), in conjunction with its receptor (cpFtsY), can target proteins co- and post-translationally to the thylakoid membrane (Gutensohn et al. 2006, Li et al. 1995). cpSRP is composed of a 54kDa GTPase (cpSRP54) subunit, homologous to the 54kDa subunit in the canonical pathway, and a 43kDa subunit (cpSRP43) unique to the chloroplast (Gutensohn et al. 2006, Li et al. 1995). In the chloroplast, there are two pools of cpSRP54 which give rise to dual functionality (Li et al. 1995). One pool of cpSRP54 associates with the ribosome and co-translationally targets proteins to their destination, while the other pool of cpSRP54 binds to cpSRP43, forming the cpSRP heterodimeric complex, and post-translationally targets lightharvesting chlorophyll-binding proteins (LHCPs) to the thylakoid membrane (Gutensohn et al. 2006, Li et al. 1995). LHCPs function as an antenna complex to focus photons for the production of high energy electrons (Büchel 2015, Pietrzykowska et al. 2014, Kong et al. 2016). LHCPs account for almost half of the membrane proteins in the thylakoids and are the most abundant protein on earth (Henry 2010, Xu et al. 2011, Xia et al. 2012). cpSRP43 evolved in the chloroplast to aid in the translocation of LHCPs without requiring the use of the ribosome or an RNA component (Schuenemann et al. 1998). cpSRP binds to LHCPs to form a soluble “transit complex” thereby enabling the LHCPs to traverse the stroma without becoming involved in inappropriate interactions (Schuenemann et al. 1998). cpSRP43 has been shown to act as a disaggregase for hydrophobic LHCPs (Falk and Sinning 2010). The aggregation of LHCPs was monitored in the presence of each component of the cpSRP pathway and cpSRP43 was shown to exclusively prevent the aggregation of LHCPs (Falk and Sinning 2010). Unlike the Hsp104/ClpB family of chaperones, cpSRP43 does not require ATPase activity for disaggregation of LHCPs (Falk and Sinning 2010). However, its ability to dissolve aggregates of LHCPs is comparable to that family of disaggregases (Jaru-Ampornpan et al. 2010, Nguyen et al. 2013, Jaru-Ampornpan et al. 2013, Falk and Sinning 2010). Studies have also revealed that Hsp70, Hsp60, and trigger factor (TF), a bacterial chaperone involved with hydrophobic regions of proteins, cannot substitute for cpSRP43 in the cpSRP pathway (Jaru-Ampornpan et al. 2010, Yuan et al. 1993). cpSRP43 interacts with the L18 region on LHCP, which is a hydrophilic peptide connecting the second and third transmembrane domains of LHCP (TM2, TM3) (Jonas-Straube et al. 2001, Tu et al. 2000). Hydrophobic interactions between the transmembrane regions of LHCP and cpSRP43 are also considered to contribute significantly to the ability of cpSRP43 to successfully chaperone the LHCPs during translocation (Jaru-Ampornpan et al. 2010, Cain et al. 2011, DeLille et al. 2000). The contribution of hydrophobic interactions is based on studies in which binding was maintained under high salt conditions (Jaru-Ampornpan et al. 2010, Cain et al. 2011, DeLille et al. 2000). In addition, 60-fold higher binding affinity between the full length LHCP *versus* the L18 peptide alone was found, revealing the involvement of TM3 of LHCP for efficient binding (Jaru-Ampornpan et al. 2010, Cain et al. 2011, DeLille et al. 2000).

cpSRP43 is a multidomain protein containing three chromodomains (CD1-3) and four ankyrin repeats (Ank1-4) located between CD1 and CD2 (Jonas-Straube et al. 2001, Schuenemann et al. 1998) (Figure 1A). While chromodomains of other proteins are known to regulate the structure of chromatin due to their ability to promote protein-protein interactions, cpSRP43 is the only non-nuclear chromoprotein (Gutensohn et al. 2006). The chromodomains interact with the methionine-rich domain of cpSRP54, resulting in the formation of heterodimeric cpSRP (Jonas-Straube et al. 2001).The crystal structure of cpSRP43 reveals two hydrophobic grooves which are separated by a positively charged ridge on one side and a highly negatively charged surface on the other side (Stengel et al. 2008). Examination of the crystal structure of cpSRP43 suggests that the charge distribution of cpSRP43 is reminiscent of the canonical SRP RNA component which acts as a scaffold for positioning the M-domain of cpSRP54 in the formation of cpSRP (Stengel et al. 2008, Batey et al. 2000). Ankyrin repeat regions are found in several different classes of proteins with varied functions, suggesting that the versatility of ankyrin repeat structural units mediates different types of protein-protein interactions (Kohl et al. 2003). Ank2 and Ank3 of cpSRP43 display the typical helix-turn-helix motif of an ankyrin domain (Stengel et al. 2008). However, the flanking Ank regions, 1 and 4, contain elongated helices (Stengel et al. 2008). The helices in Ank4 are extended by 16 residues, a noteworthy divergence from the typical Ank structure, which indicates a role in protein-protein interaction (Stengel et al. 2008). The Ank regions of cpSRP43 bind the L18 domain of LHCP (Jonas-Straube et al. 2001). The association of cpSRP43 with cpSRP54 and LHCP constitutes the soluble “transit complex” necessary for the translocation and integration of LHCP (Schuenemann et al. 1998). In addition, cpSRP43 binds to the c-terminal portion of the integral membrane protein Albino3 (Alb3) to initiate and expedite docking at the receptor for integration (Falk et al. 2010a). Reconstruction of the global structure of cpSRP43, based on small-angle X-ray scattering (SAXS) data combined with molecular dynamics (MD) simulations, reveals an elongated shape (~120 angstroms in length) which supports the proposition that this protein requires an extensive surface area for binding its substrate and cpSRP54 (Jaru-Ampornpan et al. 2010). It has also been demonstrated that cpSRP43 displays significant interdomain dynamics in studies utilizing single molecule FRET (smFRET), MD simulations and isothermal titration calorimetry (ITC) (Gao et al. 2015). Based on these studies, cpSRP43 is thought to have closed, open and extended conformational states which indicates a high degree of flexibility across the entire protein (Gao et al. 2015). Decreased flexibility upon binding to cpSRP54 was detected in conjunction with an increased affinity for the L18 region of the LHCP binding substrate, which validates the assertion that cpSRP43 undergoes changes in flexibility to execute its role in the cpSRP pathway (Gao et al. 2015).

**Figure 1.**
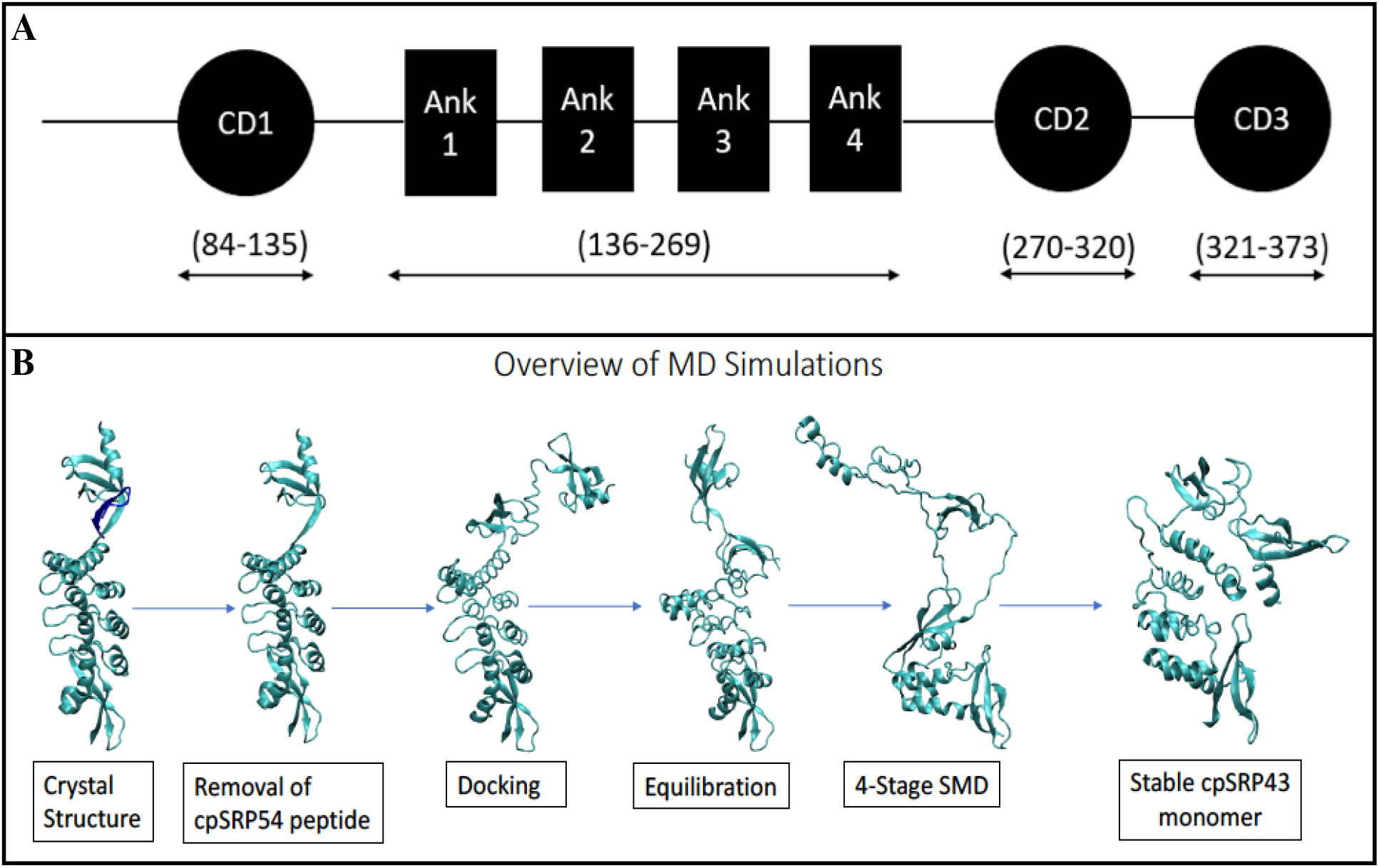
**A) Schematic diagram of the structural domains in cpSRP43.** The residues spanning each domain are indicated with double header arrows. **B) An overview of the MD simulation process.** The cpSRP54 peptide was removed from the crystal structure and the CD3 domain was docked to it. Equilibrium MD simulations of the docked model were then followed by a 4-stage SMD process. The equilibrium simulations of the 4th SMD model were extended to the microsecond level in order to further investigate a potentially stable conformation.

While computational studies of various chloroplast proteins such as the stromal ridge complex and the cpFtsY receptor have been performed (Zhang and Ding 2016, Yang et al. 2011), very little information exists on the structural and conformational dynamics of cpSRP43 or the cpSRP43/cpSRP54 complex. Recently, Gao et al reported that cpSRP43 displays significant interdomain dynamics using nanosecond-level all-atom MD simulations in conjunction with small angle x-ray scattering (SAXS) data (Gao et al. 2015). In the present study, we investigate the structural and conformational dynamics of the complete cpSRP43 structure, using a combination of equilibrium and non-equilibrium all-atom MD simulations carried out at the microsecond level.

Our simulations have revealed that cpSRP43 adopts a stable globular conformation that is substantially different from the linear crystal structures reported previously (Stengel et al. 2008, Holdermann et al. 2012). The results of these microsecond-level MD simulations clearly suggest that cpSRP43 has a stable structure containing regions that exhibit significant backbone dynamics. This conformation of cpSRP43 identified *in silico* is consistent with the results of biochemical and biophysical experiments.

## Materials and Methods

### Expression and Purification of Recombinant cpSRP43

For expression of cpSRP43, BL-21 Star cells containing pGEX-6P-2-cpSRP43 were grown to an Optical Density of 0.6–8.0 at Abs_600_ and incubated with 1 mM isopropyl β-D-thiogalactoside for 3.5 hours. Cells were harvested and resuspended in lysis buffer. Cells were sonicated for 25 cycles with 10 second on/off pulses. Supernatant was separated from the cell debris using centrifugation at 19000 rpm for 30 minutes and subsequently passed over glutathione Sepharose that was then washed extensively with equilibration buffer (2.7mM KCl, 1.8mM KH_2_PO_4_, 15mM Na_2_HPO_4_, 137mM NaCl, pH 7.2). GST-cpSRP43 was eluted using 10 mM L-Glutathione then exchanged into cleavage buffer (50 mM Tris-HCl, 150 mM NaCl, 1 mM EDTA, 1 mM dithiothreitol, pH 7.0). Overnight cleavage in solution was setup at 4°C on a rocker with 10 units of PreScission Protease per liter of original cells for 16 hours. The cleavage product was passed back onto glutathione Sepharose to separate out the GST tag and the cleaved cpSRP43 was further purified by gel filtration chromatography. Purity of proteins was visualized using 15% Sodium Dodecyl Sulfate Poly Acrylamide Gel Electrophoresis (SDS-PAGE) followed by staining with brilliant blue.

### Circular Dichroism

Circular dichroism (CD) measurements were performed on a Jasco J-1500 CD spectrometer equipped with a variable temperature cell holder. Conformational changes in the secondary structure of cpSRP43 were monitored in the Far-UV region between 190 to 250 nm using a protein concentration of 8.5 μM in 0.27 mM KCl, 0.18 mM KH_2_PO_4_, 1.5 mM Na_2_HPO_4_, 13.7 mM NaCl (pH 7.2) in a 1 mm pathlength quartz cuvette. The scan speed, band width and data pitch were set to 50nm/min, 1.00nm and 0.1nm, respectively. Three scans were collected (within a 1000 HT voltage range) and averaged to obtain the CD spectra. The thermal denaturation/renaturation scans were recorded from 25 to 80°C at 5°C increments.

### Fluorescence Spectrometry

All fluorescence measurements were performed on a Hitachi F-2500 spectrophotometer at 25 °C using a slit width of 2.5 nm and a 10 mm quartz cuvette. Intrinsic fluorescence experiments were performed with protein concentration of 0.2 mg/mL in in 0.27 mM KCl, 0.18 mM KH_2_PO_4_, 1.5 mM Na_2_HPO_4_, 13.7 mM NaCl (pH 7.2) All samples were excited at a wavelength of 280 nm and the emission spectra were recorded from 300 nm to 500 nm.

### Equilibrium Unfolding

Thermal denaturation of cpSRP43 as probed by circular dichroism was performed on a Jasco-1500 spectrophotometer using a protein concentration of 8.5 μM in 0.27mM KCl, 0.18mM KH_2_PO_4_, 1.5mM Na_2_HPO_4_, 13.7mM NaCl (pH 7.2). Spectra were collected at 5-degree increments from 25 to 90 °C. Molar ellipticity values were recorded and plotted as a function of temperature. Thermal denaturation as probed by fluorescence was carried by manual heating of samples at 5-degree increments for 5 minutes.

### Limited Trypsin Digestion

Limited trypsin digestion of cpSRP43 and Ovalbumin was performed in phosphate buffer containing 2.7mM KCl, 1.8mM KH_2_PO_4_, 15mM Na_2_HPO_4_, 137mM NaCl (pH 7.2). The initial reaction tube contained 8.5 μM of protein and 0.5 μg of enzyme. The trypsin-containing samples were incubated at room temperature (25 °C). Digested samples were removed every 2 minutes for up to 16 minutes. The reaction was stopped by the addition of 10% trichloroacetic acid at the end of 20 minutes after which the samples were resolved on a 15% sodium dodecyl sulfate-polyacrylamide gel electrophoresis (SDS-PAGE) gel and subsequently stained using Coomassie Blue. UN-ScanIT software (Silk Scientific Inc.) was applied to identify the percent of cpSRP43 digested at each time point. Ovalbumin (A5503) was purchased from Sigma Co., St. Louis.

### Fluorescence Quenching

All measurements were made using Fluorescence Spectrophotometer F-2500 (Hitachi). The excitation wavelength was set at 295 nm and data was collected in the range of 315-450 nm. Fluorescence emission intensities were recorded for cpSRP43 at 340 nm upon titration with stock solutions of acrylamide, cesium chloride, potassium iodide and succinimide. Analysis of the quenching data was made using the Stern Volmer equation [1] and the modified Stern-Volmer equation [2] (Eftink and Ghiron 1981).

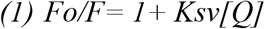

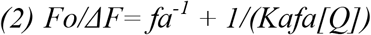

Where *Fo* and *F* are the relative fluorescence intensities in the absence and presence of each quencher, respectively and [Q] is the concentration of each quencher. *Ksv* is the Stern-Volmer quenching constant for each quencher, *fa* is the fraction of fluorescence accessible to each quencher and *Kd* is the corresponding quenching constant.

### Equilibrium MD Simulations of cpSRP43 and cpSRP43/cpSRP54

We set out to determine the stability of cpSRP43 compared to the cpSRP43/cpSRP54 complex. We used NAMD 2.11-13 (Phillips et al. 2005) and CHARMM36m all-atom additive force field (Huang et al. 2017) parameters to perform equilibrium MD simulations based on crystal structures obtained from the RCSB Protein Data Bank (Berman et al., 2000). We based these simulations on an X-ray crystal structure containing both cpSRP43 and cpSRP54 in complex (PDB entry: 3UI2) (Holdermann et al. 2012). This allowed us to simulate both cpSRP43 and the cpSRP43/cpSRP54 complex. This was accomplished by manually removing the cpSRP54 sequence from the PDB file. Both systems were solvated in a TIP3P water box of size 150 Å X 150 Å X 150 Å and neutralized with 0.15 M NaCl. Both the cpSRP43 (96358 atoms) and cpSRP43/cpSRP54 (96390 atoms) systems were energy-minimized for 10,000 steps using the conjugate gradient algorithm (Reid 1971). They were then relaxed by applying harmonic restraints (force constant = 1 kcal/molÅ^2^) to the C_α_ atoms for 1 ns. The initial relaxation was performed in an NVT ensemble while production runs were performed in an NPT ensemble. Simulations were carried out using a 2-fs time step at 300 K using a Langevin integrator with a damping coefficient of γ = 0.5 ps^-1^. The pressure was maintained at 1 atm using the Nosé-Hoover Langevin piston method (Reid 1971, Martyna et al.1994). The smoothed cutoff distance for non-bonded interactions was set to 10-12 Å and long-range electrostatic interactions were computed with the particle mesh Ewald (PME) method (Darden et al. 1993). A production run of 100 ns was carried out for each system.

### Molecular Docking of CD3

To model the complete cpSRP43 structure, we used VMD (Humphrey et al. 1996) tcl scripting to combine two different crystal structures – cpSRP43 (without the CD3 domain) (PDB:3UI2) Holdermann et al. 2012) and the CD3 domain (PDB:2N88) (Horn et al. 2015). We used the Sequence Alignment tool in VMD (Humphrey et al. 1996) to find a common amino acid for docking. GLY 316 was chosen as the anchor point for the docking procedure as it was shared by both structures. The script used the common amino acid to anchor the two PDBs and generate the docked model. The final structure had the following amino acid sequence - CD1(85-135), Ank1-Ank4(136-269), CD2 (270-320), CD3 (321-369), cpSRP54(528-540) (see Figure S1).

### Equilibrium MD Simulations of Docked Model (with and without cpSRP54)

The docked model was solvated in a 170 Å X 170 Å X 170 Å TIP3P water box and neutralized with 0.15 M NaCl. The systems were minimized and relaxed using the same procedure as described above for the initial set of simulations. The cpSRP43 and the cpSRP43/cpSRP54 systems had approximately 130800 and 130945 atoms, respectively. A production run of 100 ns was carried out for each system as described above for the initial set of simulations (see Figure S1).

### Non-equilibrium SMD simulations

Starting from the equilibrated model of the docked cpSRP43 (without cpSRP54), we used the collective variable module (Fiorin et al. 2013) in NAMD (Phillips et al. 2005) to perform non-equilibrium steered MD (SMD) simulations based on the previously published SAXS data (Jaru-Ampornpan et al. 2010, Gao et al. 2015). The Sequence Alignment tool in VMD (Humphrey et al. 1996) was used to determine the common residues between our SMD model (equilibrated docked model) and the SAXS conformations (40 frames based on variations of 4 different conformations). The alignment produced common residues from all major functional domains. The residues from Ank1-4, CD2, and CD3 were used to identify the most similar SAXS conformation to the initial SMD model (SAXS Model 1). Once the first target conformation was determined, non-equilibrium SMD simulations were used to steer the equilibrated docked model towards the target SAXS conformation. Two separate distance vectors were used in each simulation as collective variables - Dist_(cD2,Ank1-4)_ and Dist_(cD3,Ank1-4)_. A distance vector collective variable is defined based on the vector connecting the mass center of a particular domain to the mass center of another domain. In our case, both collective variables shared one of their domain selections, i.e., the Ankyrin repeats (136-269). Each distance vector also had a unique domain selection, with Dist_(CD2,Ank1-4)_ having CD2 (270-320) (determining its position with respect to Ankyrin 1-4) and Dist_(CD3,Ank1-4)_ having CD3 (321-369) (determining its position with respect to Ankyrin 1-4). The mass center of the three domains from the SAXS model were used as target centers for the docked model. The SMD simulation was run for 10 ns to generate SMD Model 1. A force constant of 500 kcal/mol Å^2^ was used for the SMD simulation. The final conformation of this simulation was used as the initial conformation for the second SMD simulation to generate SMD Model 2. A similar procedure as the first SMD simulation was used to find the most similar SAXS target conformation (SAXS Model 2) and perform a SMD simulation similar to the first SMD simulation. This process was then repeated for SMD/SAXS Model 3 and SMD/SAXS Model 4 (see Figure S1).

### Equilibrium MD Simulations of SAXS Models and SMD Generated Models

The final conformation of each SMD simulation (SMD Models 1-4) was used as the initial conformation for a 100 ns equilibrium simulation. The production runs were performed using the same protocol described for the initial set of equilibrium simulations. The simulation for SMD Model 4 was extended to 200 ns.

SAXS Models 1-4 were solvated in a 170 Å X 170 Å X 170 Å TIP3P water box and neutralized with 0.15 M NaCl. The systems were minimized and relaxed using the same protocol described above for the initial set of equilibrium simulations. Production runs were performed for 100 ns each (see Figure S1).

### Microsecond-level All-Atom Equilibrium MD Simulation of SMD Model 4

The equilibrium MD production run for the 4^th^ SMD model was extended for 2 microseconds on the Anton2 supercomputer (Shaw et al. 2014), with a timestep of 2.5 fs. The pressure was maintained at 1 atm isotropically, using the MTK barostat, while the temperature was maintained at 300 K, using the Nosé–Hoover thermostat. The long-range electrostatic interactions were computed using the fast Fourier transform (FFT) method (Young et al. 2009). Conformations were collected every 240 picoseconds.

The Root Mean Square Deviation (RMSD) of Cα atoms were calculated using VMD (Humphrey et al. 1996). Forward RMSD was calculated with respect to the starting conformation of a trajectory. Reverse RMSD was calculated with respect to the final conformation of a trajectory. Root Mean Square Fluctuation (RMSF) of individual residues was calculated based on the Cα atoms using VMD. The salt bridge identification and distance measurements were done using the salt bridge plugin in VMD (Humphrey et al. 1996). Strong salt bridges were defined as interactions with a donor-acceptor cut-off distance below 4 angstroms for at least 40% of the total simulation time. Solvent accessible surface area (SASA) was calculated using the internal SASA measurement method of VMD (Humphrey et al. 1996).

## Results and Discussion

Starting from the 3UI2 crystal structure (Holdermann et al. 2012), a stable conformation of monomeric cpSRP43 was identified using microsecond-level equilibrium MD simulations. The cpSRP54 peptide was removed from the crystal structure, after which the CD3 domain was docked to the crystal structure. This was followed by equilibrium MD simulations of the docked structure for 100 ns. SMD was then performed in 4 stages based on different SAXS conformations to generate 4 SMD models. The 4^th^ SMD model was found to be stable and was thus selected for further investigation. A 2 microsecond equilibrium-MD simulation of the 4^th^ SMD model then produced a conformation of monomeric cpSRP43 that was stable for approximately 1.6 microseconds. An illustrated flow chart of the procedure is shown in Figure 1B.

### cpSRP54 Stabilizes cpSRP43

Initially, we performed 100 ns equilibrium MD simulations with both cpSRP43 and the cpSRP43/cpSRP54 complex to determine their relative structural stability. RMSF analysis (Figure S2A) showed that the CD2 domain is the most flexible region of cpSRP43 and that fluctuations in this region decrease significantly when cpSRP54 is present. This is corroborated by RMSD analysis (Figure S2B), which showed that cpSRP43 is more stable when complexed with cpSRP54. The results of these analyses thus indicate that cpSRP54 stabilizes cpSRP43 when it interacts with the CD2 domain. A unique salt bridge between the CD2 domain of cpSRP43 and cpSRP54 possibly contributes to the stability of the SRP43/SRP54, but more experimentation would be required to determine its importance.

### The CD3 Domain Introduces Structural Instability to both cpSRP43 and the cpSRP43/cpSRP54 Complex

We performed 100 ns equilibrium MD simulations to determine whether the molecular docking of the CD3 domain to cpSRP43 introduced structural instability. RMSF analysis showed that there was an increase in fluctuation throughout the cpSRP43 structure for both the cpSRP43 and cpSRP43/cpSRP54 systems (Figure S3A, S3C). In addition, RMSD analysis also showed that the introduction of the CD3 domain clearly resulted in the destabilization of cpSRP43 from both systems (Figure S3B, S3D). Salt bridge analysis did not reveal any significant differences between the systems with CD3 and without CD3, thus indicating that the introduction of the CD3 domain contributed directly to the differential behavior displayed by the systems with CD3.

### Comparative Analysis of the Stability of SMD-generated cpSRP43 Models versus Similar SAXS Models

An accumulated work profile from our SMD simulations (Figure S4A) revealed that there was a surge in work output of around 500 kcal/mol for the Dist_(CD2,Ank1-4)_ collective variable, which was then followed by a drop in work output of around 200 kcal/mol. This indicates that a transition state was potentially crossed during the generation of SMD Model 4. Equilibrium simulations based on the SAXS models and their corresponding SMD generated models (see Methods), were then performed in order to investigate the structural and conformational dynamics of each SAXS and SMD model. A total of 8 simulations were performed, based on 4 SAXS models and 4 corresponding SMD models. Comparison of forward and reverse RMSDs (see Methods) showed that SAXS and SMD Models 1-3 were relatively unstable (Figure S4-S6). On the other hand, the reverse RMSD of SMD Model 4 showed that it was moving towards a stable conformation (Figure S5).

### Identifying a Stable Conformation for Monomeric cpSRP43 using Microsecond-level MD Simulations

As SMD Model 4 showed a stable conformation, its equilibrium production run was extended by 2 microseconds. It was believed that this starting conformation would remain stable throughout the simulation. However, several fluctuations were observed during the first 0.4 microseconds. The system then settled down into a more stable conformation which was maintained for approximately 1.6 microseconds (Figure 2A-B). The CD2 and CD3 domains are the most flexible regions of this stable conformation (Figure 2C-D). This is consistent with our previous results, which indicated that the docking of CD3 to the crystal structure caused a decrease in the stability of cpSRP43. A visual inspection of the trajectory revealed that ankyrin repeats 2 and 3 were rotated by 180 degrees in the final conformation, when compared to the crystal structure. To quantify this structural change, we calculated the combined solvent accessible surface area (SASA) of all tryptophan residues as well as the SASA for individual tryptophan residues. The average SASA of all tryptophan residues was 445 Å^2^ for the crystal structure and 565 Å^2^ for the stable conformation. The SASA for the individual tryptophan residues is shown in Table 1. Five out of eight tryptophan residues in the stable monomer show an increase in average SASA. Our data thus indicates that there is a fairly significant increase in the average SASA for all tryptophan residues in the stable conformation.

**Figure 2.**
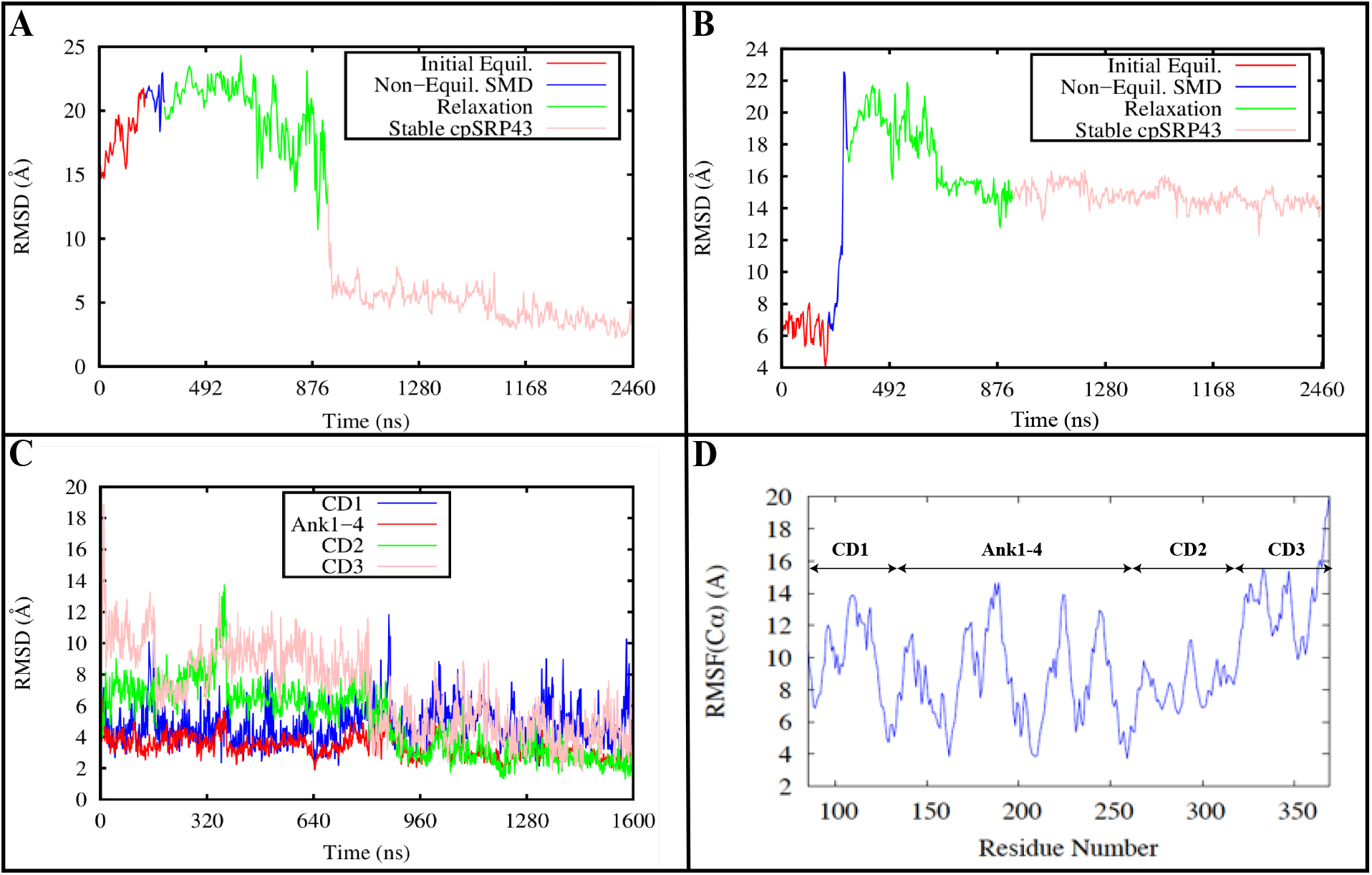
Microsecond-level MD simulations reveal a stable conformation of monomeric cpSRP43. A) Reverse RMSD plot of monomeric cpSRP43. B) Forward RMSD plot of monomeric cpSRP43. The RMSD plots show that cpsRP43 settles into a stable conformation after 0.4 microseconds and remains stable for 1.6 microseconds. C) RMSD and D) RMSF plots of individual cpSRP43 domains show that the most flexible regions of the stable monomeric cpSRP43 are the CD2 and CD3 domains. This is consistent with the results obtained when CD3 was docked to the crystal structure.

**Table 1.** A comparison of the average SASA of individual tryptophan residues. There is a net increase in SASA for the tryptophan residues in the stable monomer compared to the corresponding residues in the crystal structure.

| **Tryptophan residue number** | **Average SASA – crystal structure (Å<sup>2</sup>)** | **Average SASA – stable monomer (Å<sup>2</sup>)** |
| --- | --- | --- |
| 106 | 62 | 95 |
| 114 | 112 | 107 |
| 132 | 2 | 27 |
| 133 | 13 | 55 |
| 291 | 16 | 47 |
| 299 | 83 | 88 |
| 343 | 52 | 43 |
| 351 | 104 | 101 |

In addition to this, the beta strand connecting CD2 and ankyrin repeat 4 in the crystal structure, becomes a collapsible loop in the stable monomer. These structural changes facilitate the interaction of CD1 with ankyrin-4 and CD3 (Figure 3), thus enabling different domains of the protein to come closer to each other than in the crystal structures (Stengel et al. 2008, Holdermann et al. 2012). Our data indicates that the average distance between CD1-CD2 and CD2-ankyrin changes significantly (Table 2). The interaction of these domains results in the protein adopting a more globular structure as opposed to the mostly linear crystal structures (Stengel et al. 2008, Holdermann et al. 2012).

**Figure 3.**
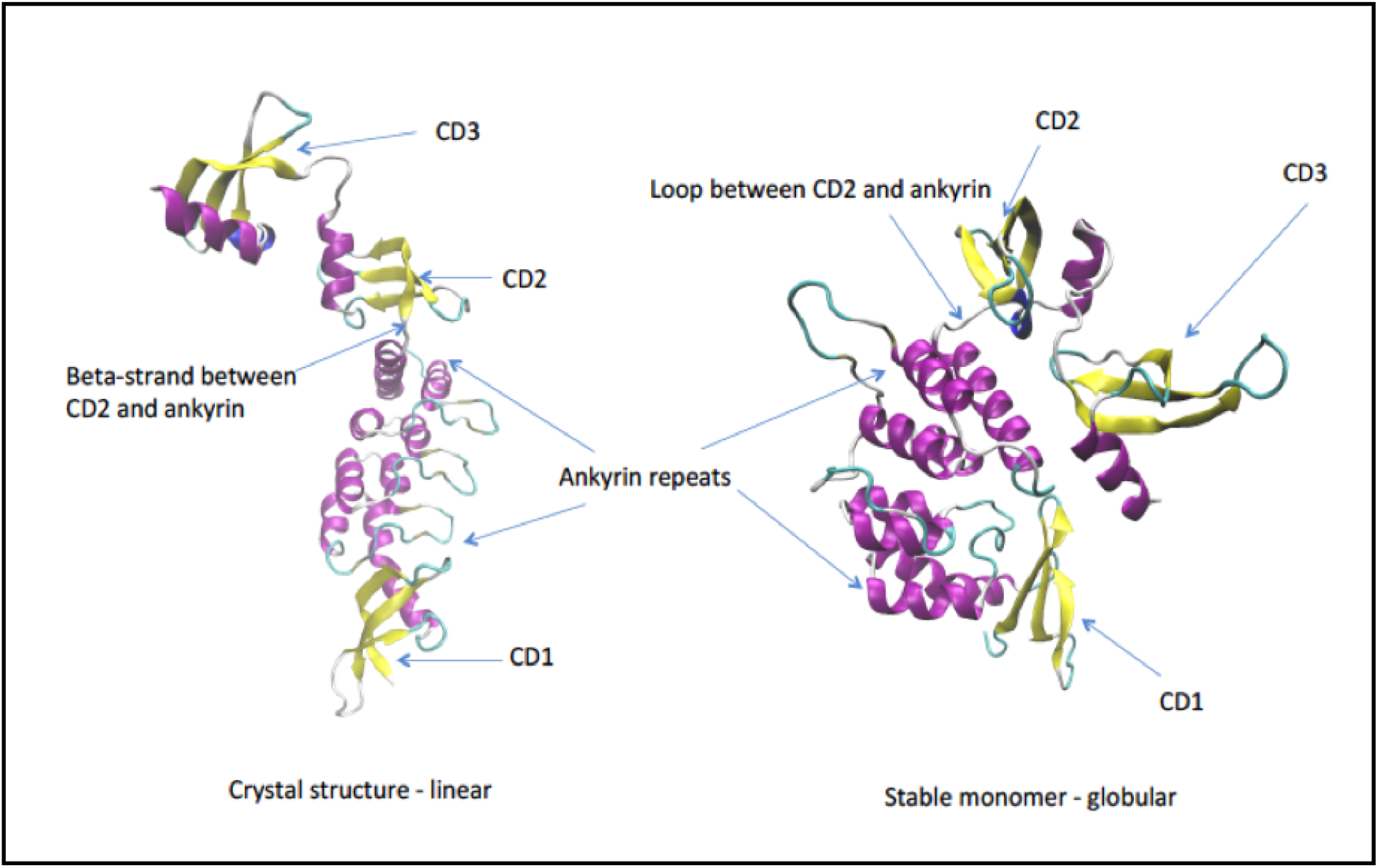
Stable monomeric cpSRP43 adopts a globular conformation. A 180-degree rotation of the middle ankyrin repeats coupled with the transformation of a beta strand to a loop (between CD2 and Ank1) causes different domains to interact. This results in the stable monomer adopting a globular conformation as opposed to the linear conformation of the crystal structure.

**Table 2.** A comparison of the average distance between domains for the crystal structure and the stable monomer. There is a significant decrease in interdomain distance for CD1-CD2 and CD2-Ank in the stable monomer. A much smaller reduction in interdomain distance occurs for CD1-Ank.

| **Domain pairs** | **Average distance in crystal structure** | **Average distance in stable monomer** |
| --- | --- | --- |
| CD1-CD2 | 62.74 | 49.54 |
| CD1-Ank | 29.46 | 27.74 |
| CD2-Ank | 35.99 | 29.80 |

To further investigate the dynamics of this conformation, a comparative analysis of the salt-bridges present in the stable monomer and the crystal structure was performed. Interestingly, seven unique interdomain salt-bridges were found in the stable monomeric cpSRP43, while no interdomain salt-bridges were present in the crystal structure (Holdermann et al. 2012) (Figure 4A-G). An illustration of each interdomain salt bridge is shown in Figure 4H. Out of these seven salt-bridges, one was formed between CD1 and Ank1-4, three between Ank1-4 and CD2, two between CD2 and CD3 and one between Ank1-4 and CD3. In terms of presence during the simulation, the salt bridges GLU129-ARG161 (78% - CD1 to Ank1-4), GLU 256-LYS283 (62%) and GLU298-ARG226 (54%) (both Ank1-4 to CD2) are particularly significant. A total of 21 salt bridges were found in the stable cpSRP43 monomer, with 7 novel interdomain salt bridges, 9 novel intradomain salt bridges and 5 salt bridges shared with the crystal structure (Table S1). A total of 15 salt-bridges were found in the crystal structure (Table S2).

**Figure 4.**
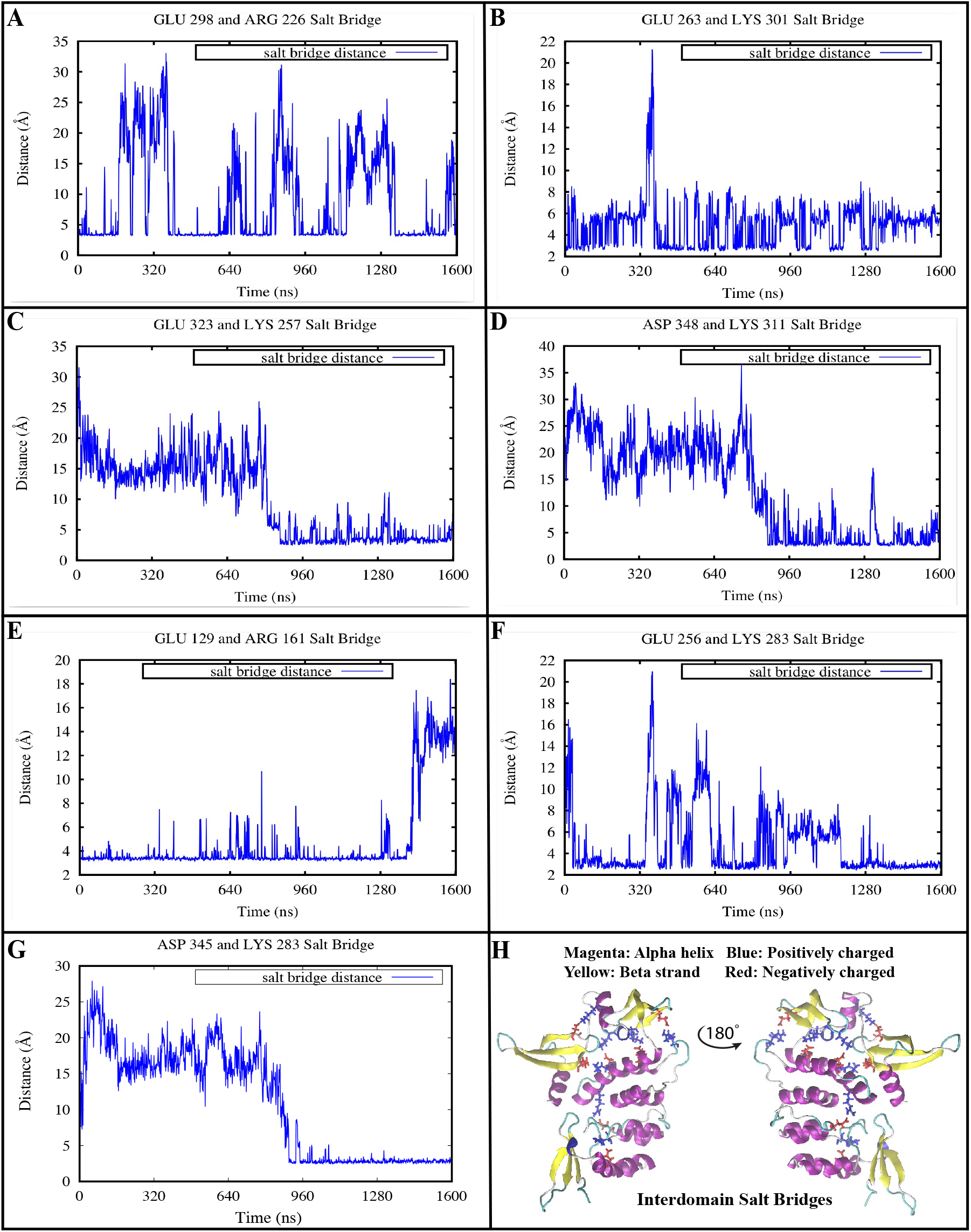
Seven interdomain salt bridges occur in the stable cpSRP43 monomer but not in the crystal structure. A) GLU298-ARG226 (Ank1-4 and CD2); B) GLU263-LYS301 (Ank1-4 and CD2); C) GLU323-LYS257 (Ank1-4 and CD3); D) ASP348-LYS311 (CD2 and CD3); E) GLU129-ARG161 (CD1 and Ank1-4); F) GLU256-LYS283 (Ank1-4 and CD2); G) ASP345-LYS283 (CD2 and CD3); H) Cartoon representation of the unique interdomain salt-bridges that occur in the stable monomer.

Our data clearly shows that the 180-degree rotation of the ankyrin repeats coupled with other secondary structural changes effectively accounts for the presence of unique interdomain saltbridges in the stable cpSRP43 monomer. The conformational change forced various domains to interact with each other, thus resulting in the formation of stable interdomain salt-bridges. These salt bridges reinforced the overall effects of the conformational change, thus maintaining the stable conformation for 1.6 microseconds.

### Secondary Structure Analysis using Far-UV-CD Spectroscopy

Far UV-circular dichroism [190-250nm] is a useful tool to probe the secondary structure of proteins in native conformations) and to evaluate any changes in conformation upon alterations made to the microenvironment of the protein. The far UV CD spectrum of cpSRP43 shows a double negative minimum at 208nm and 222nm suggesting that the backbone of the protein is predominately folded into a helix conformation. This aspect is consistent with the crystal structure of cpSRP43 (Figure 5A) (Stengel et al. 2008, Holdermann et al. 2012, Horn et al. 2015, Sivaraja et al. 2005, Kathir et al. 2008). Triple resonance NMR data for the CD1 domain of cpSRP43 revealed that its backbone is predominantly folded into a beta-barrel structure (Sivaraja et al. 2005). NMR data for the CD2 and CD3 domains suggests that they also fold predominantly into a beta-sheet conformation with helical segments at their C-terminal ends (Sivaraja et al. 2005). Therefore, the CD spectrum of intact cpSRP43 appears to stem largely from the Ank domains which are helical.

**Figure 5.**
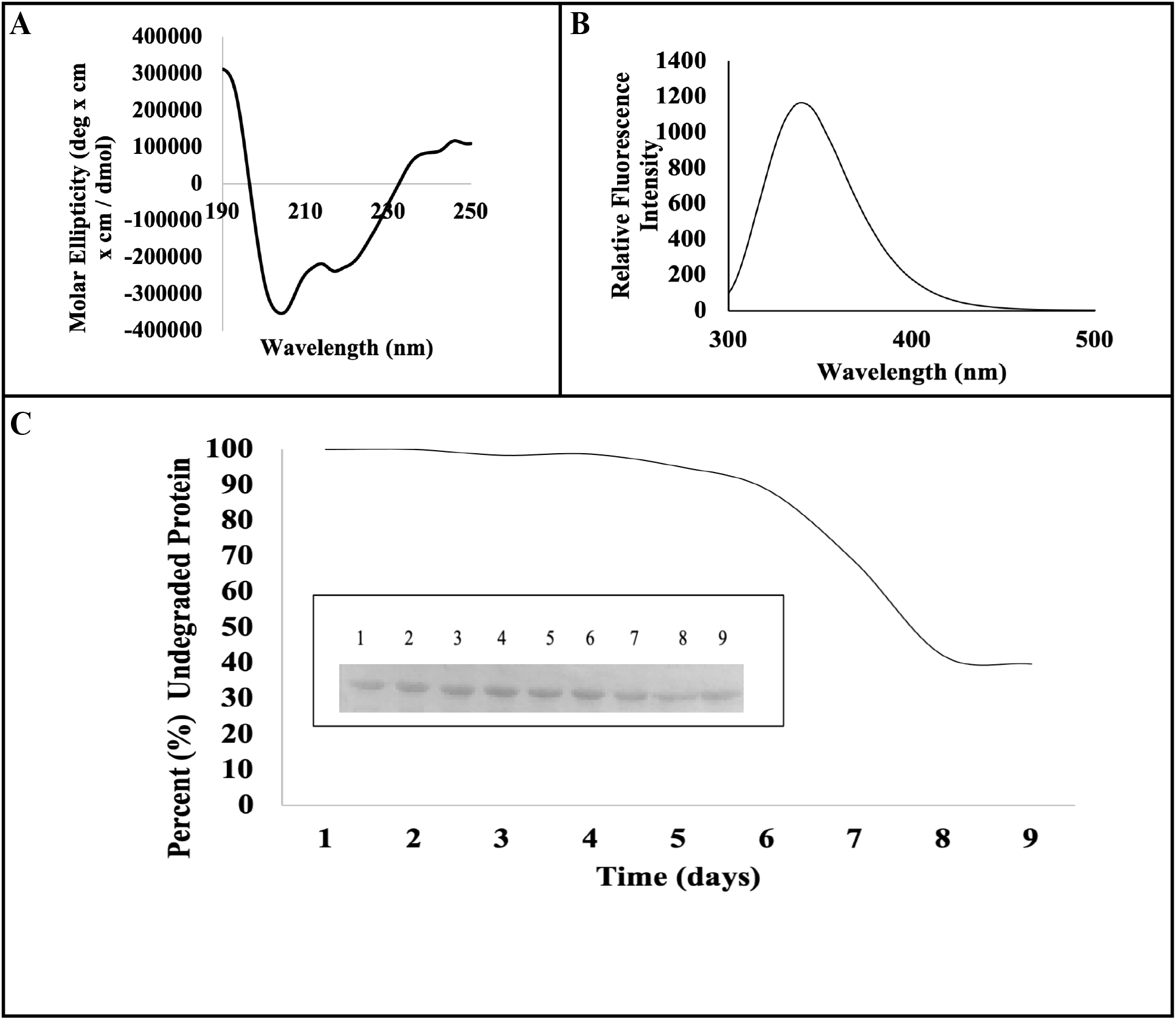
A) Far-UV CD and B) Intrinsic Fluorescence spectra of cpSRP43. C) Densitometric plot depicting the rate of degradation of cpSRP43, incubated at 25° C, over a nine-day period. The insert shows a portion of the 15% SDS-PAGE with lanes, 1-9, representing days 1-9.

### Monitoring of Tertiary Structural Change by Fluorescence

Intrinsic protein fluorescence is sensitive to the local environment surrounding the indole ring of tryptophan units and is useful in determining the stability of a protein by tracking the folding/unfolding rates under different conditions (Royer 1995). The intrinsic fluorescence spectrum of cpSRP43 in its native conformation shows an emission maximum at 340nm (Figure 5B). cpSRP43 contains 8 tryptophan residues and all are located in the three chromodomains. CD1 contains four tryptophan residues. CD2 and CD3 contain two tryptophan residues each. The wavelength of maximum emission of cpSRP43 indicates that most of the tryptophan moieties in the protein are in a partially buried microenvironment.

### Protein Stability and Degradation at Room Temperature

Storage of proteins at room temperature often leads to their degradation. In many cases, proteins cannot withstand being left at room temperature for more than a day whereas some proteins may degrade almost immediately. Protein stability is governed by a balance of different forces. The rate of degradation of cpSRP43 was monitored at 25 °C, over a nine-day period (Figure 5C). We found that cpSRP43 was almost completely resistant to degradation over four days. In fact, insignificant degradation was observed even after seven days at room temperature. The average pixels recorded for the SDS-PAGE gel (samples for days 1-9) using the UN-SCAN It densitometric software are as follows: 8.9,8.9,8.75,8.78,8.47,7.9,6.09,3.75 and 3.54, respectively.

### Thermal Stability of cpSRP43

Thermal denaturation and renaturation of cpSRP43 was probed by circular dichroism from 25-90° C at 5° C increments. The Far UV CD spectra of cpSRP43 acquired in the temperature range of 25°C to 90°C are nearly identical and superimpose well suggesting that the protein is stable in this temperature range (Figure 6A). The extraordinary thermal stability of cpSRP43 is interesting considering its molecular mass and significant internal dynamics. CD spectra of cpSRP43 heated to 90°C and subsequently cooled back to 25 °C showed that the thermal unfolding of the protein is completely reversible. The extraordinary structural stability and reversibility of the thermal unfolding are characteristic features of heat shock proteins that exhibit protein folding chaperone activity. Interestingly, as mentioned earlier, cpSRP43 is known to facilitate the disaggregation of LHCP (Falk and Sinning 2010). cpSRP43 retained its secondary structure after heating and cooling cycles which proves to be an interesting feature for such a structurally dynamic protein. The individual circular dichroism spectra taken at each temperature increment show negligible change in comparison to one another after heating and cooling cycles (Figure 6B). The spectra of cpSRP43 before heating (25° C), upon heating to 90° C and after cooling back down to 25° C appear nearly identical when superimposed (Figure 6C). The conformity between spectra is also illustrated by the ratio of the double negative minima at each temperature increment (Figure 6A-B inset). Upon heating, cpSRP43 shows a steady decrease in intrinsic fluorescence intensity (Figure 6D). Intrinsic fluorescence readings taken of cpSRP43 before heating show that the protein is in the native conformation with an emission maximum at 340nm. The samples were then heated to 90° C at which point the emission maxima reached 348nm and the relative intensities diminished, indicating that the protein was in an unfolded state. The fluorescence readings were taken again after the protein cooled down to room temperature and the emission maxima returned to 340nm suggesting that, upon cooling, the protein was able to regain its native fold (Figure 6D). Thermal stability data reveals that cpSRP43 is highly stable at extreme temperatures. The structure of cpSRP43 contributed by the backbone arrangement is not perturbed under high heat conditions and although the tertiary structure may undergo some changes upon heat exposure, any shift in the folded state of cpSRP43 is reversible upon cooling.

**Figure 6.**
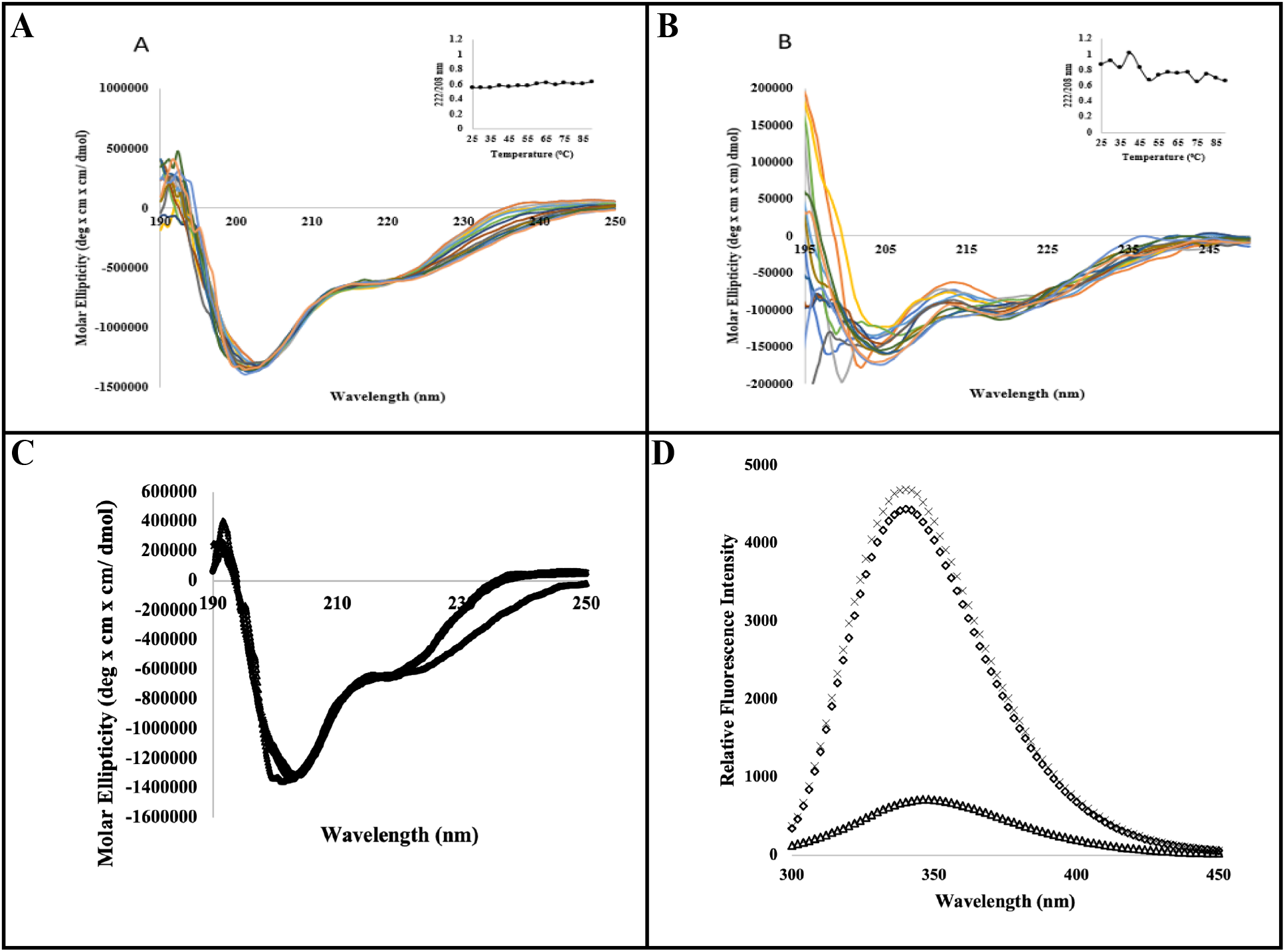
**A) Thermal denaturation and B) renaturation of cpSRP43 as probed by circular dichroism at five-degree increments from 25 −90 degrees celsius.** Ratio plots (222/208 nm) for each spectra overlay are depicted in the inset graphs. **C) Thermal stability of cpSRP43 monitored by far UV-CD spectroscopy. D) Thermal stability of cpSRP43 monitored by intrinsic fluorescence spectroscopy.** Legend - Native cpSRP43 at 25° C (o), cpSRP43 heated to 90° C (Δ) and cpSRP43 upon cooling back down to 25° C (x).

### Susceptibility of cpSRP43 to Trypsin Digestion

Limited trypsin digestion (LTD) provides useful information on the conformational flexibility of a protein (Olsen et al. 2004). In this context, we evaluated the backbone flexibility of cpSRP43 by LTD. The rate of trypsin-mediated digestion of the protein was assessed based on the intensity of the ~ 35 kDa band corresponding to uncleaved cpSRP43. In addition, LTD of ovalbumin (molecular mass of ~ 45 kDa) was carried out. Comparison of the two was used to evaluate the conformational flexibility of cpSRP43. As shown in Figure 7A-B, cpSRP43 was highly susceptible to proteolytic cleavage which is reflected by the need to carry out the experiment at 25° C in the presence of a small quantity of trypsin. In initial trials in which cpSRP43 was subjected to digestion at varying concentrations of trypsin at 37° C, the degradation occurred almost immediately which necessitated the optimization of the experimental parameters to slow down the trypsin induced digestion. cpSRP43 was easily digested within twenty minutes whereas ovalbumin effectively remained undigested under similar conditions. The resistance of ovalbumin to trypsin is surprising considering the fact that it contains 33 potential trypsin cleavage sites as opposed to 29 trypsin cleavage sites in cpRSP43. These results suggest that the backbone of cpSRP43 is highly flexible despite its extraordinary thermal stability. The high backbone flexibility discerned from the results of the LTD experiments is very consistent with the structure of cpSRP43 characterized by SAXS, smFRET, and MD simulation data. (Liang et al. 2016, Gao et al. 2015).

**Figure 7.**
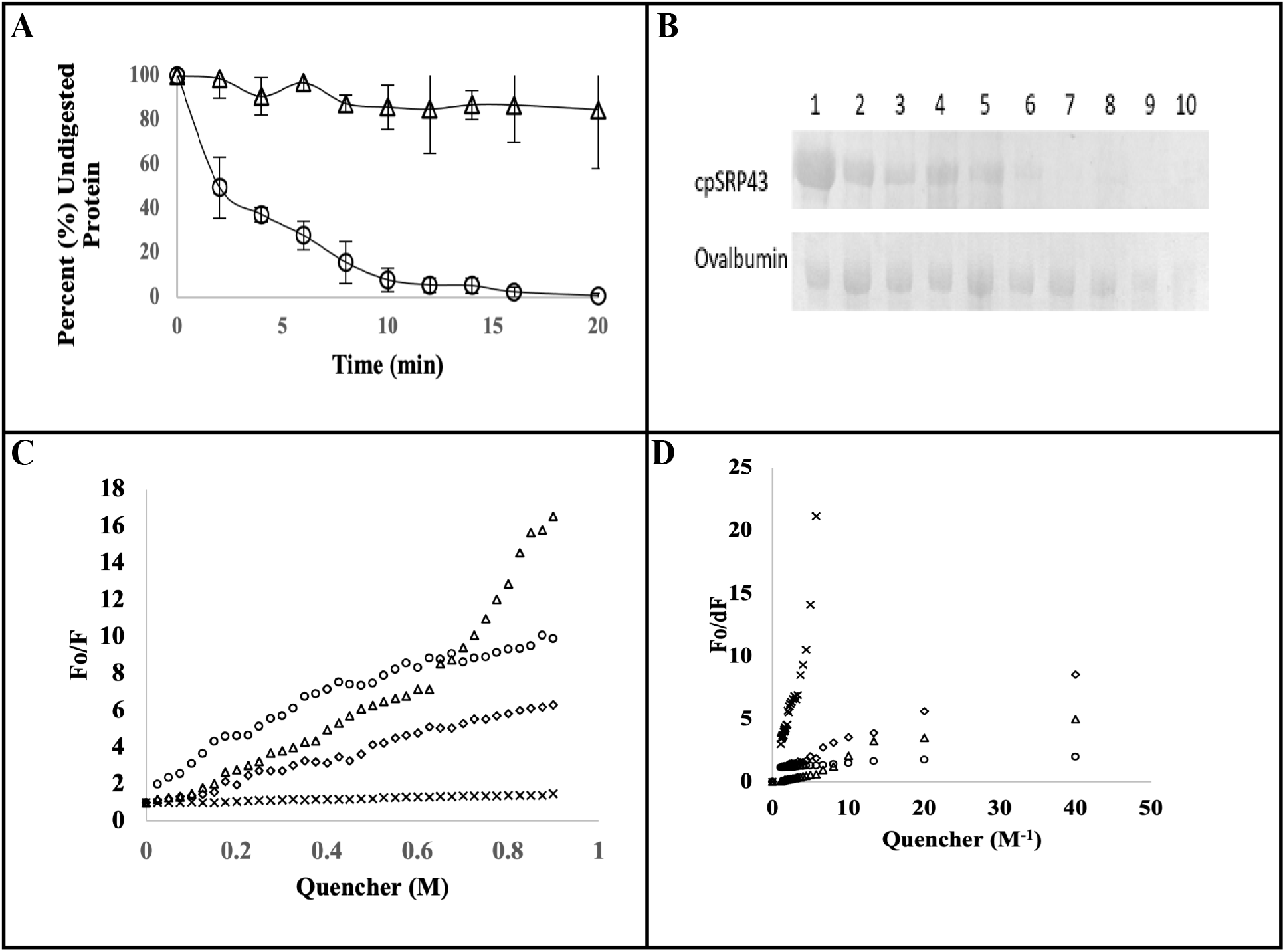
**A) Densitometric analysis of the time dependent limited trypsin digestion of cpSRP43 (o) and Ovalbumin (Δ) as monitored by SDS-PAGE.** B) Coomassie blue stained bands of cpSRP43 and Ovalbumin. **C) Stern-Volmer plots and D) Modified Stern-Volmer plots of fluorescence quenching of cpSRP43.** Legend - Acrylamide (Δ), Cesium Chloride (x), Succinimide (⟡) and Potassium Iodide (o).

### Fluorescence Quenching of cpSRP43

The intrinsic fluorescence of tryptophan is sensitive to changes in the polarity of the medium and therefore can provide useful information on the tertiary structure of proteins (Di Giambattista et al. 1984). Therefore, monitoring the microenvironment of tryptophan residues can facilitates the understanding of subtle conformational changes in proteins. The accessibility of tryptophan moieties within a protein can be monitored through fluorescence quenching (Di Giambattista et al. 1984, Samworth et al. 1988, Jamir and Seshagirirao 2018, Papadopoulou et al. 2005). Acrylamide and succinimide are both neutral quenchers and can potentially quench tryptophan residues located in different microenvironments. However, as succinimide is a bulkier molecule, it cannot access the tryptophan residues located in the interior of proteins. Succinimide can only access the superficially and partially buried tryptophan residues (Di Giambattista et al. 1984). Anionic (I^-)^ and cationic (Cs^+^) quenchers were also used to obtain information on the microenvironment of the tryptophan residues. Quenching accessibility of ionic quenchers depends on the charged environment in the spatial vicinity of the tryptophan residues (Di Giambattista et al. 1984). Fluorescence quenching parameters were obtained from Stern-Volmer and modified Stern-Volmer plots (Figure 7C-D). All Stern-Volmer plots showed a linear upward curvature except for the cesium chloride plot which was a flat line in comparison to the other quenchers. The Ksv values of acrylamide, potassium iodide and succinimide are significantly larger than that of cesium chloride which indicates that they are more efficient quenchers for cpSRP43 (Table 3). Acrylamide and succinimide displayed high fluorophore accessibility suggesting that the tryptophan residues are largely exposed to solvent throughout the protein. Accessibility was higher for potassium iodide than cesium chloride, which indicates that the local environment of tryptophan residues within cpSRP43 is predominantly positively charged. Given the high accessibility of these tryptophan residues to a range of different fluorophores and the relatively high SASA values of these residues (both individual and combined) from the MD simulations (Table 1), these collective results are in agreement with our observations of a stable conformation that shows significant inter-domain dynamics and backbone flexibility.

**Table 3:** Fluorescence quenching of cpSRP43 using different quenchers. Ksv: Stern-Volmer quenching constant, Kd: quenching constant, fa: fraction of the total fluorescence accessible to a given quencher.

| Quencher | K<sub>sv</sub> (M<sup>-1</sup>) | K<sub>d</sub> (M<sup>-1</sup>) | fa |
| --- | --- | --- | --- |
| Acrylamide | 10.24 | 0.82 | 0.91 |
| Succinimide | 6.0 | 0.73 | 0.89 |
| Potassium Iodide | 13.23 | 0.42 | 0.90 |
| Cesium Chloride | 0.50 | 0.99 | 0.69 |

### Comparing Experimental smFRET and Computational MD Data

Gao et al. have previously studied the structural flexibility of cpSRP43 in the presence and absence of cpSRP54 using smFRET experiments (Gao et al. 2015). Among the five constructs designed to characterize the structural dynamics of full length cpSRP43 in the presence and absence of full length cpSRP54, three showed cpSRP54-dependent behavior (see Table 4). Using our MD simulation data from both isolated cpSRP43 and the cpSRP43 in complex with the cpSRP54 fragment, we have calculated FRET distributions for the same constructs that qualitatively agree with the experimental data, providing further evidence for the validity of our computational model (see Fig. 8 for Construct 1 and Fig. S7 for Constructs 2 and 3). For the isolated cpSRP43, we used the last 1,600 ns of the 2-*μ*s equilibrium MD simulation of the docked cpSRP43, which represents the stabilized isolated cpSRP43 model proposed in this study. For the cpSRP43/cpSRP54 complex, the 100-ns production equilibrium simulation of the docked cpSRP43 in complex with the short cpSRP54 fragment discussed above was used. The methodological details concerning the construction of FRET distributions from MD simulation data are based on the study by Baucom et al. (Baucom et al. 2021).

**Figure 8.**
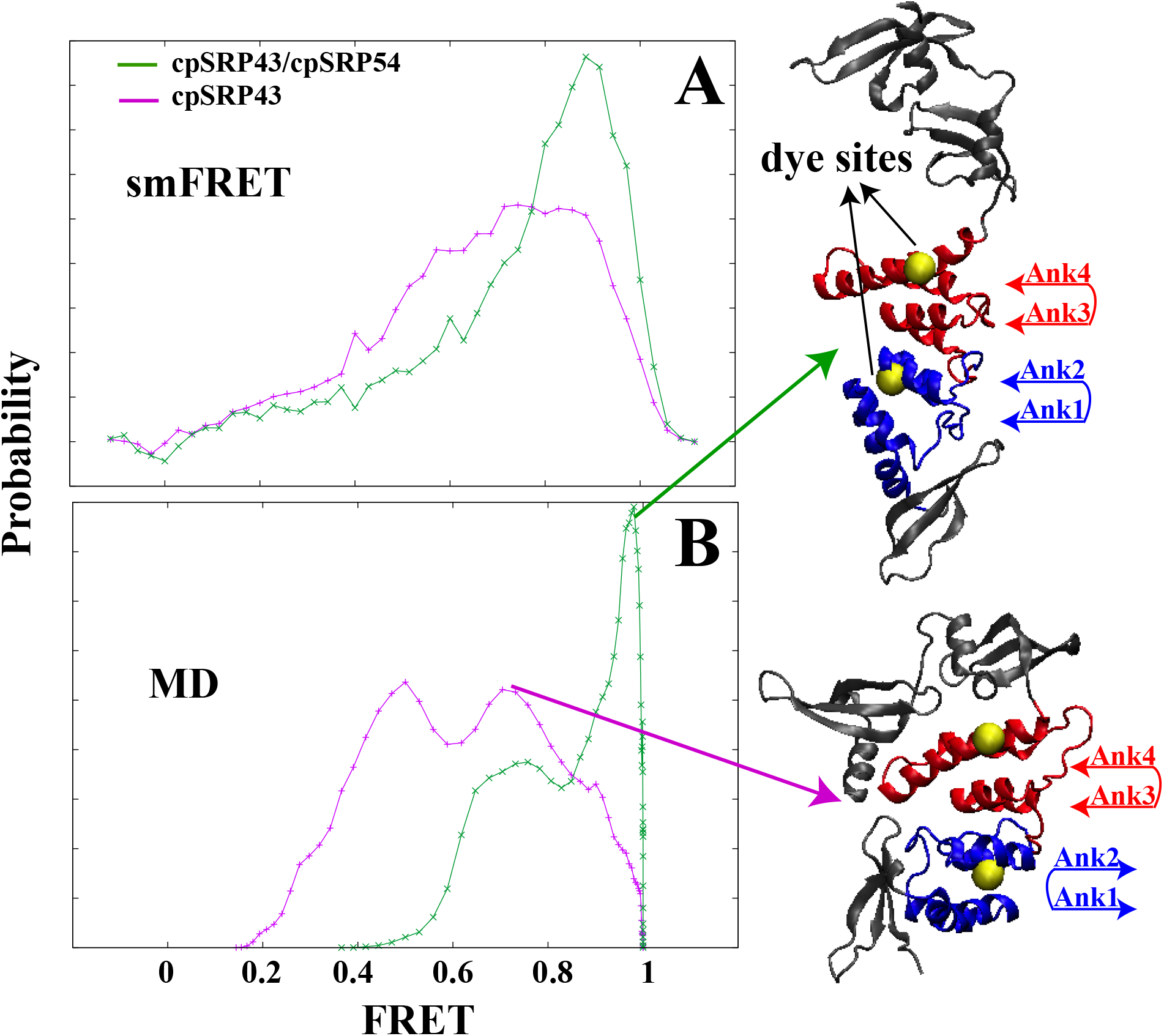
Comparing experimental smFRET (A) and computational MD data (B) for FRET distributions of a cpSRP43 mutant (Construct 1 in Table 4) in the absence (purple) and presence (green) of the cpSRP54 protein/fragment (A/B). The experimental data was previously reported in (Gao et al. 2015).

**Table 4:** Description of smFRET experiments that show differential FRET distributions for the full length cpSRP43 in the presence and absence of full length cpSRP54 (Gao et al. 2015).

| Construct | cpSRP43 Mutations | Labeled Sites | Labeled Domains |
| --- | --- | --- | --- |
| 1 | V265C, C301A | Cys-179, Cys-265 | Ank2/Ank4 |
| 2 | C179A, R333C | Cys-301, R333C | CD2/CD3 |
| 3 | None | Cys-179, Cys-301 | Ank2/CD2 |

## Conclusion

We have investigated the structure and stability of cpSRP43 under different conditions by utilizing various biophysical and *in silico* techniques. cpSRP43 is found to be a highly heat stable protein capable of maintaining its secondary structure upon subjection to significant increases in temperature. In addition, cpSRP43 was able to resist degradation when left for several days at room temperature and can refold itself after cooling cycles. The degree of thermal stability displayed by cpSRP43 would be expected of a highly compact or rigid protein. However the data derived from limited trypsin digestion experiments are indicative of a highly flexible and dynamic structure. The extraordinary stability of cpSRP43 is unusual considering the parallel degree of high flexibility and structural dynamics within this protein. These aspects were previously unexplained by the existing crystal structure(s) of cpSRP43. The dynamics of the cpSRP43 were analyzed based on all atom MD simulations, which provide a high degree of chemical and physical specificity. Our original hypothesis concerning the cpSRP43 and cpSRP43/cpSRP54 complex was confirmed based on structural fluctuation. The addition of the CD3 domain to the existing cpSRP43 crystal structure (Holdermann et al. 2012) revealed that CD3 destabilizes the protein significantly. SAXS and SMD based simulations were performed and analyzed in order to find a stable conformation of cpSRP43. SAXS data samples the predominant protein conformations in solution, thus it provided us with coordinates of several conformations which served as targets for the SMD simulations. All four SAXS models demonstrated significant instability. Although all four of our SMD models were more stable over 10 ns compared to the SAXS models, three of these failed to produce stable conformations. The accumulated work profile indicates that the first 3 SMD models had little work output, whereas SMD Model 4 showed a significant increase in work output followed by an abrupt decrease. This could potentially indicate crossing a transition state that produced the stable conformation. The methods we employed to identify a stable conformation of cpSRP43 are novel and can be applied to similar problems using a combination of SAXS and non-equilibrium steered molecular dynamics. SMD Model 4 was investigated further and found to be stable over 1.6 microseconds. This stable monomeric conformation of cpSRP43 was found to adopt a globular structure, with different interactions and secondary structural features compared to the crystal structure (Holdermann et al. 2012). Ankyrin repeats 2 and 3 undergo a 180-degree rotation and the beta strand connecting CD2 to ankyrin-4 becomes a loop. When this loop collapses, CD1 interacts with the ankyrins and CD3. Our analysis shows that the distance between the various domains decreases significantly in the stable monomer when compared to the crystal structure, accounting for the presence of seven unique inter-domain saltbridges. This combination of secondary structural changes and interactions results in the globular structure of the stable monomer which is significantly different from the linear crystal structure (Holdermann et al. 2012).

Circular dichroism and fluorescence techniques were utilized to study the thermal stability as well as the unfolding and subsequent refolding of cpSRP43. In addition, results of limited trypsin digestion of cpSRP43 show a high degree of flexibility in the backbone structure of cpSRP43 in comparison with a commensurate protein. The results of intrinsic quenching experiments of cpSRP43 are in agreement with *in silico* solvent accessibility data of tryptophan residues, providing further confirmation of the flexible nature of cpSRP43. In addition, our comparative MD simulation data on isolated cpSRP43 and the cpSRP43/cpSRP54 complex are in qualitative agreement with previously published experimental smFRET data, providing further evidence for the validity of our computational models. Therefore, using a combination of experimental techniques and microsecond-level all-atom MD simulations, we have isolated a stable conformation of monomeric cpSRP43 that displays inherent flexibility and significant dynamic behavior. Our study has thus provided new insights into the structural and conformational dynamics of cpSRP43, which may prove to be biologically relevant with respect to its function as a chaperone.

## Acknowledgments

This research was supported by the U.S. Department of Energy (grant no. DE-FG02-01ER15161), National Science Foundation grants CHE 1945465 and OAC 1940188, the National Institutes of Health/National Cancer Institute (NIH/NCI) (1 RO1 CA 172631), the NIH through the COBRE program (P30 GM103450), and the Arkansas Biosciences Institute. Anton 2 computer time was provided by the Pittsburgh Supercomputing Center (PSC) through Grant R01GM116961 from the National Institutes of Health. The Anton 2 machine at PSC was generously made available by D.E. Shaw Research. This work also used the Extreme Science and Engineering Discovery Environment (allocation MCB150129), which is supported by National Science Foundation grant number ACI-1548562.

## Notes

### Competing Interest Statement

The authors have declared no competing interest.

