## Supporting Information for "cpSRP43 is both highly flexible and stable: Structural insights using a combined experimental and computational approach"

**Mitchell Benton, Mercede Furr, Vivek Govind Kumar, Feng Gao, Colin David Heyes, Thallapuranam Krishnaswamy Suresh Kumar* and Mahmoud Moradi*.**
Department of Chemistry and Biochemistry, University of Arkansas
Fayetteville, AR, 72701, USA

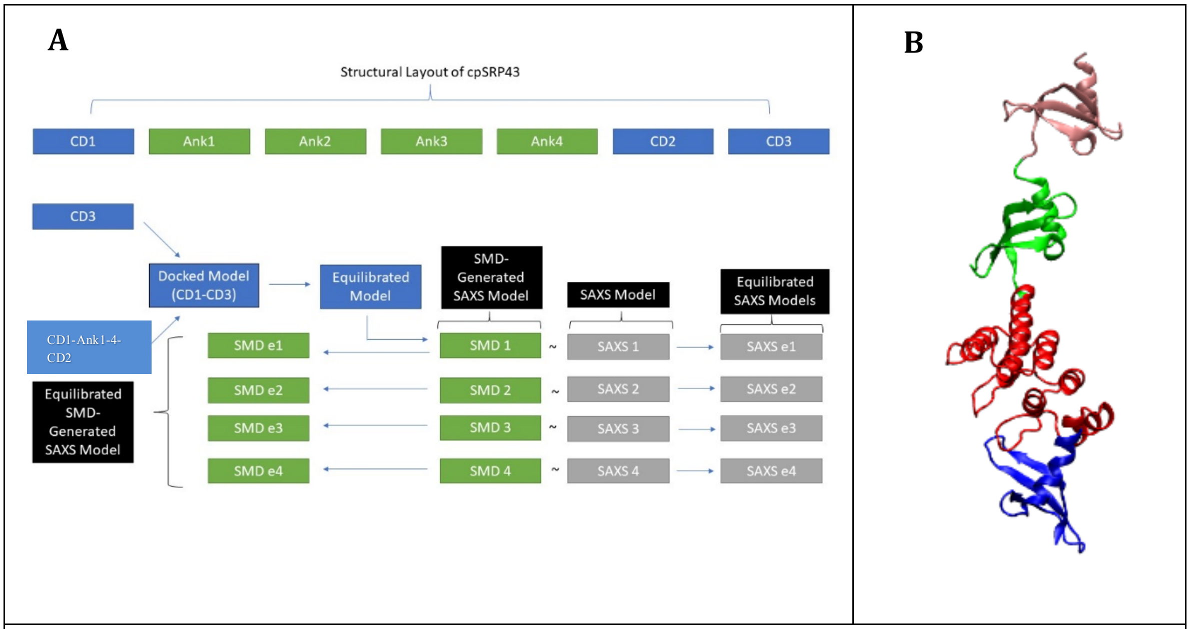


**Figure S1. SMD-SAXS simulation flowchart. A)** A flowchart outlining the simulation procedure used to generate SMD models from our docked cpSRP43 model is presented here. The docked model was generated from two different PDBs and was then equilibrated. An initial SAXS model was used in conjunction with the equilibrated docked model to generate SMD Model 1. This process was repeated until SMD Model 4 was generated. See Methods section for more details. **B)** Cartoon representation of the complete structure of cpSRP43, with CD1(blue), Ank1-4(red), CD2(green), and CD3(pink) domains.

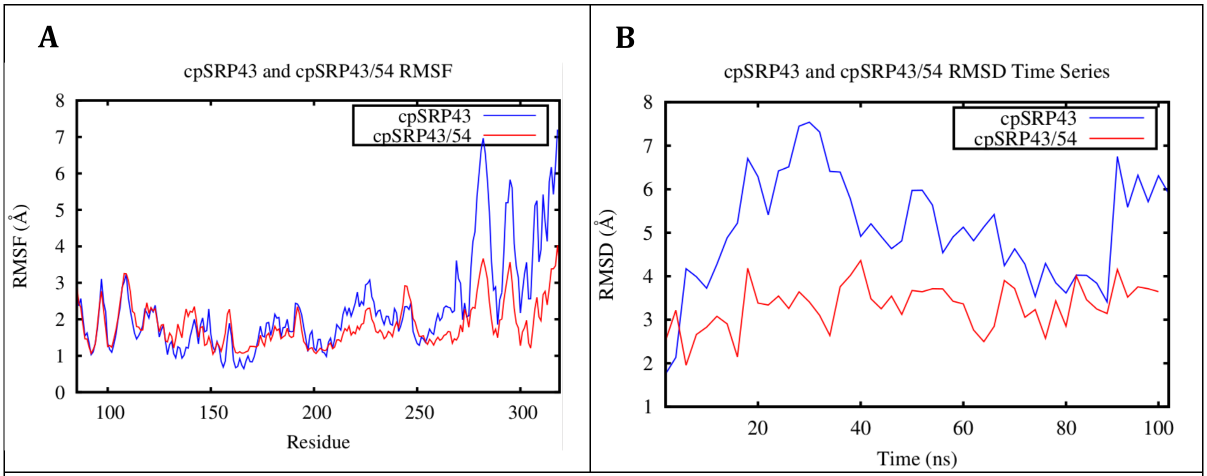


**Figure S2. cpSRP54 stabilizes cpSRP43. A)** RMSF analysis of cpSRP43 from the monomeric cpSRP43 (blue) and cpSRP43/54 (red) systems shows that the CD2 domain is the most flexible part of the protein. Fluctuations in this region are reduced significantly when interacting with cpSRP54. **B)** RMSD analysis clearly shows that cpSRP43 is more stable in the presence of cpSRP54. (Initial equilibrium simulations before docking).

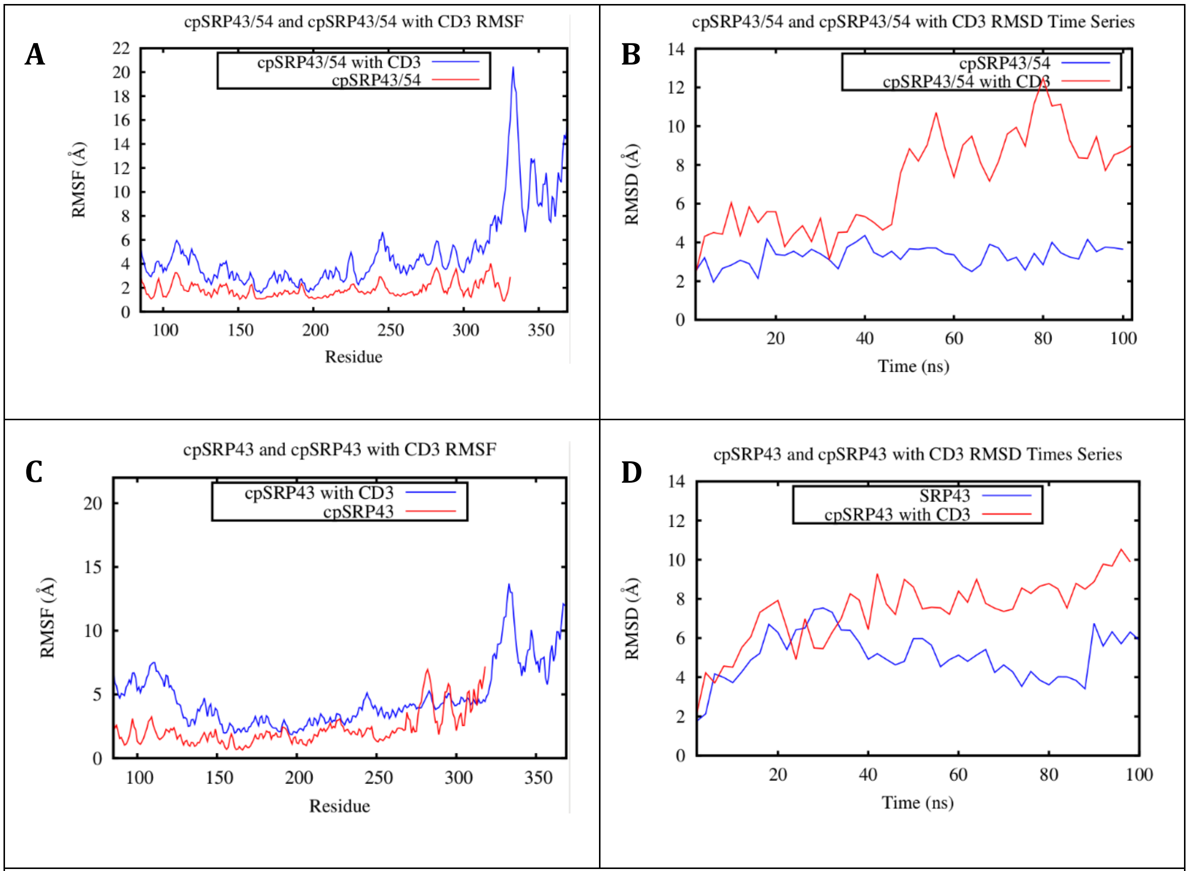


**Figure S3. The CD3 domain destabilizes cpSRP43 in the presence and absence of cpSRP54. A & C)** RMSF analysis clearly showed that there was an increase in fluctuation for all regions of the cpSRP43 structure, in both the monomeric cpSRP43 and cpSRP43/cpSRP54 trajectories. **B & D)** RMSD analysis also showed that the introduction of the CD3 domain clearly resulted in the destabilization of cpSRP43 from both systems. (Equilibrium simulations of CD3-docked models).

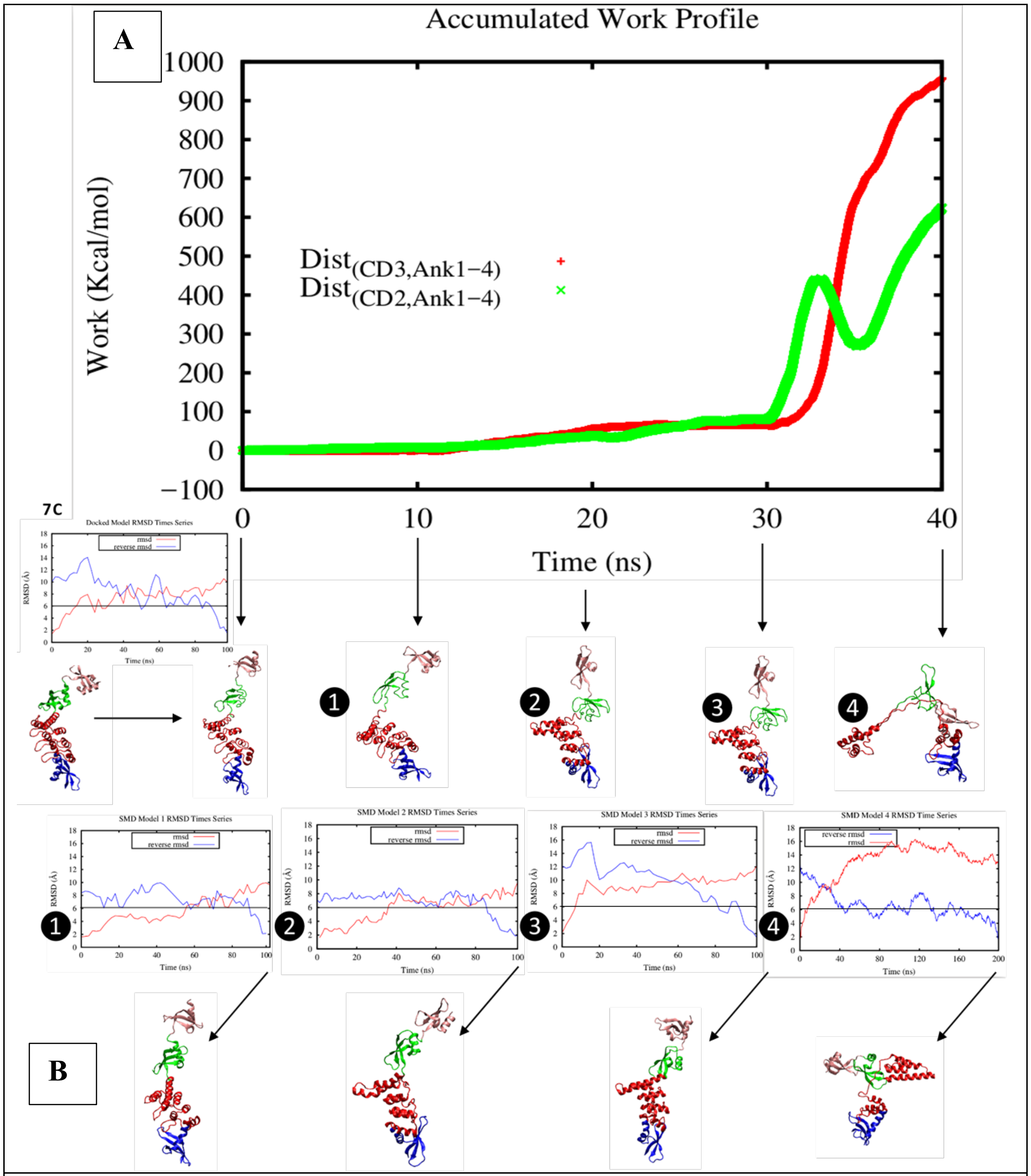


**Figure S4. Stability of SMD-generated cpSRP43 models. A)** This figure illustrates the accumulated work output and associated conformations from the SMD simulations. The conformation at 0 ns represents the final conformation from the equilibrium simulation of the docked model. The final conformations of SMD simulations 1-4 were generated at 10 ns intervals and are illustrated below the work profile. **B)** The numbers next to the forward/reverse RMSD plots represent the initial conformations used for equilibrium simulations based on the final SMD conformations 1-3 (100 ns) and 4 (200 ns). SMD model 4 is relatively stable compared to the other models. **C)** Forward and reverse RMSD time series of the docked model from a 100 ns equilibrium simulation (inset).

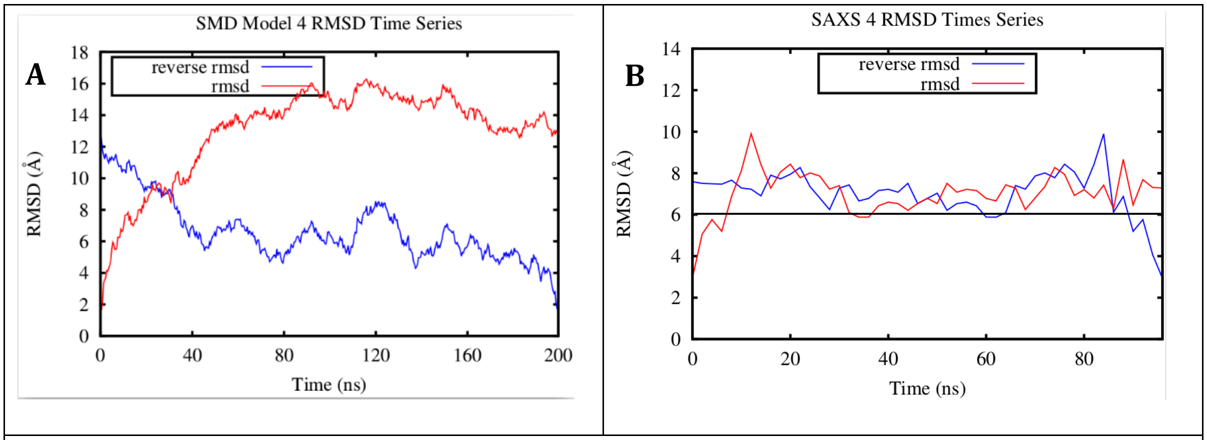


**Figure S5. Comparison of SMD/SAXS Model 4. A)** SMD Model 4 appears to be relatively unstable based on forward RMSD, but the reverse RMSD plot suggests that it is actually fairly stable over 200 ns. Microsecond-level equilibrium simulations of this model were then performed. **B)** The corresponding SAXS model is relatively unstable overall in comparison to the SMD model.

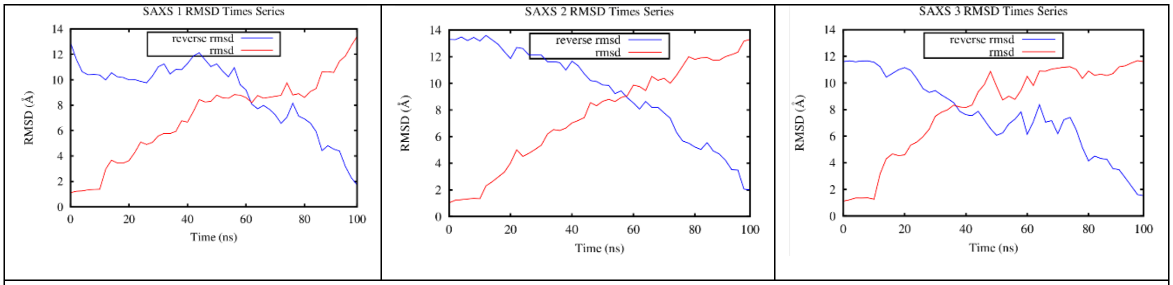


**Figure S6. Forward and reverse RMSD time series for SAXS models 1-3.**

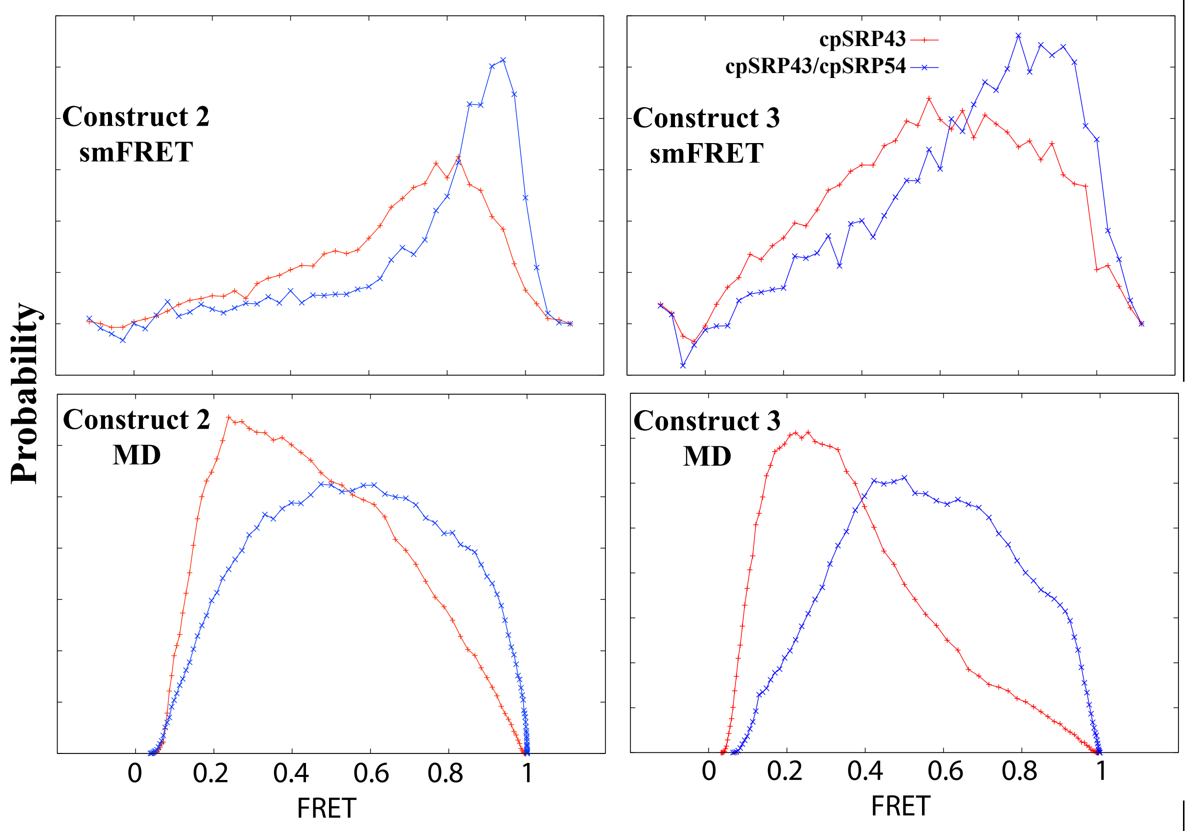


**Figure S7. Comparing FRET distributions of cpSRP43 (red) and cpSRP43/cpSRP54 (blue) from smFRET experiments (top) and MD simulations (bottom) for Constructs 2 and 3 (see Table 4). The experimental data was previously reported in Gao et al. 2015. Also see Figure 8 for the same data for Construct 1. In all cases, there is a shift towards higher FRET values when cpSRP54 (in the experimental case) or cpSRP54 fragment (in the computational case) is present.**

| Salt bridge | Occupancy (%) |
| --- | --- |
| E263-R235 | 97 |
| E323-K342 | 95 |
| E338-R329 | 84 |
| E286-R279 | 81 |
| E129-R161 | 78 |
| E318-K281 | 75 |
| E101-R93 | 66 |
| E318-K278 | 65 |
| E256-K283 | 62 |
| E127-K88 | 56 |
| E298-R226 | 54 |
| D154-R251 | 52 |
| D345-K283 | 41 |
| D273-K292 | 39 |
| E263-K301 | 38 |
| E323-K257 | 36 |
| E314-K278 | 33 |
| E181-R251 | 33 |
| D348-K311 | 31 |
| E286-K301 | 31 |

**Table S1. Important salt bridges in stable monomeric cpSRP43**. Red = salt bridges shared by stable monomeric cpSRP43 and crystal structure (3UI2); Green = novel interdomain salt bridges; Blue = novel intradomain salt bridges.

| Salt bridge |
| --- |
| D157-R137 |
| D224-R226 |
| D273-R290 |
| D315-K311 |
| E101-R93 |
| E105-K88 |
| E141-K174 |
| E158-R137 |
| E211-K257 |
| E215-R177 |
| E256-R252 |
| E263-R235 |
| E286-R279 |
| E286-K301 |
| E314-K278 |

**Table S2. Salt bridges in the crystal structure**. Red = salt bridges shared by stable monomeric cpSRP43 and crystal structure (3UI2); Blue = novel intradomain salt bridges. The 3UI2 crystal structure contained no interdomain salt bridges.
